# Benchmark and integration of resources for the estimation of human transcription factor activities

**DOI:** 10.1101/337915

**Authors:** Luz Garcia-Alonso, Mahmoud M Ibrahim, Denes Turei, Julio Saez-Rodriguez

**Affiliations:** European Molecular Biology Laboratory - European Bioinformatics Institute (EMBL-EBI), Wellcome Genome Campus, CB10 1SD Cambridge, UK.; OpenTargets,Wellcome Genome Campus, CB10 1SD Cambridge, UK.; Joint Research Centre for Computational Biomedicine (JRC-COMBINE); RWTH Aachen University, Faculty of Medicine, Pauwelsstrasse 19, 52074 Aachen, Germany.; Department of Nephrology; RWTH Aachen University, Faculty of Medicine, Pauwelsstrasse 30, 52074 Aachen, Germany.

**Author notes:** Current address: Institute of Computational Biomedicine, Heidelberg University, Faculty of Medicine, Bioquant - Im Neuenheimer Feld 267, 69120 Heidelberg, Germany (Julio Saez-Rodriguez).

**Keywords:** Transcription Factor, Regulatory networks, Protein activities

## Abstract

Prediction of transcription factor (TF) activities from the gene expression of their targets (i.e. TF regulon) is becoming a widely-used approach to characterize the functional status of transcriptional regulatory circuits. Several strategies and datasets have been proposed to link the target genes likely regulated by a TF, each one providing a different level of evidence. The most established ones are: (i) manually curated repositories, (ii) interactions derived from ChIP-seq binding data, (iii) *in silico* prediction of TF binding on gene promoters, and (iv) reverse-engineered regulons from large gene expression datasets. However, it is not known how these different sources of regulons affect the TF activity estimations, and thereby downstream analysis and interpretation. Here we compared the accuracy and biases of these strategies to define human TF regulons by means of their ability to predict changes in TF activities in three reference benchmark datasets. We assembled a collection of TF-target interactions among 1,541 TFs, and evaluated how the different molecular and regulatory properties of the TFs, such as the DNA-binding domain, specificities or mode of interaction with the chromatin, affect the predictions of TF activity changes. We assessed their coverage and found little overlap on the regulons derived from each strategy and better performance by literature-curated information followed by ChIP-seq data. We provide an integrated resource of all TF-target interactions derived through these strategies with a confidence score, as a resource for enhanced prediction of TF activities.

## INTRODUCTION

Regulation of gene expression programs is fundamental for cell development, differentiation and tissue homeostasis. Dysregulation of such programs is responsible for most aberrant cell phenotypes, including cancer and other complex diseases. Due to their ability to interact with specific DNA regulatory regions, transcription factors (TF) are the key proteins in gene-specific transcriptional regulation, linking it to the signaling transduction network. Consequently, TFs have been proposed as downstream readouts of pathway activities and the assessment of their activity status has gained much attention in the last years. Noteworthy examples are their use in the characterization of driver somatic mutations and the identification of new markers of drug response in cancer^1–4^, or the reconstruction of the regulatory processes dictating cell differentiation^5–7^.

With high-throughput measurements of TF activities not available, a common practice is to estimate them computationally, from the gene expression levels of their direct targets (the so-called TF regulon). The assumption behind is that the level of protein activity of a TF is reflected on the transcript levels of its targeted genes^8^. Accurate TF activity quantifications will, therefore, depend on the availability of high confidence sets of functional targets, where the TF has a direct regulatory effect on the transcription of the target gene, and the specificity of the TF-target interaction, so that the regulation of the target’s transcription can be unambiguously assigned to the TF. Moreover, if the TF activity quantification approach is intended to be generally applicable to any cell type, context-independent TF-target interactions are preferred so that the predictions are consistent and comparable across cell types^4,9^. This is key for systematic studies using heterogeneous populations of cell types, such as cell line panels spanning through different tumor types or differentiation stages.

Several strategies and resources have been proposed to define the set of target genes directly regulated by a TF. These can be grouped according to the strength of evidence supporting a TF-target interaction. The first types are resources collecting manually curated interactions from peer-reviewed literature. Literature-curated resources are expected to contain high-quality TF-target regulatory interactions with experimental evidence. To our knowledge, there are around a dozen of literature-curated databases collecting interactions for human^8,10–20^. However, these differ in their curation protocols, literature selection criteria or quality controls^21^. Consequently, there is a small overlap between resources that generates uncertainty on which ones should be used^20^. In addition, they have a biased coverage towards well studied TFs, in particular those involved in diseases. Another type of strategies are high-throughput measurements of TF-DNA binding such as chromatin immunoprecipitation (ChIP)^22,23^ or DNase I hypersensitivity (DNase) assays^24^ coupled to DNA sequencing (ChIP-seq and DNase-seq). These provide high-resolution maps of *in vivo* DNA-binding regions for a given TF. Still, most binding events represent non-functional interactions, meaning that TF binding does not correspond with changes in the expression levels of the targeted gene^25,26^. As for the literature-curated resources, TF-binding peaks derived through these *in vivo* methodologies are relative to the experimental conditions and cell types used, as well as maintain biases in biomedical research. To overcome part of the limitations inherent to the mentioned experimental conditions, an alternative is to computationally predict TF-target interactions making use of TF binding sites (TFBSs) models. *In silico* identification of TFBSs on gene regulatory regions relies on the assumption that TFs have binding preferences to specific DNA sequences, referred to as a “binding motifs”^27–29^. TF binding motifs, generally modelled as position weight matrices (PWM), are then used to scan the regulatory sequences in a genome to identify candidate target genes^30^. Consequently, predictions are restricted to TFs for which binding motifs are known. Additionally, these suffer from a significant number of false positives, because the binding never occurs *in vivo* or is non-functional, or because of the uncertainty on which site is regulating which gene. Finally, TF-target interactions also can be reverse-engineered *in silico* from large-scale gene expression profiles derived from a condition of interest^31,32^. The assumption is that the transcript levels of a TF are informative of the expression levels of their targeted genes. This approach overcomes several limitations of the previous TFBSs prediction method such as the cell type-specificities (i.e. predictions are tailored towards the underlying expression profile) and the TF coverage (i.e. regulons can be inferred for any TF whose expression varies sufficiently in the corresponding gene expression dataset). Still, the approach may fail to infer TF-target interactions for TFs regulated at a molecular level other than transcription (such as post-translational modification and protein-protein interactions)^33^ and their power to distinguish direct and indirect regulation is controversial^34–36^. Taken together, currently there is no universal strategy to identify all *bona fide* targets of the full collection of TFs across all possible cell conditions.

Despite their broad use, there is no systematic evaluation of the impact of the evidence supporting the TF-target interactions in the estimation of TF activities. This is important, as they can affect substantially the results and thereby downstream analysis and interpretation. To address this, we performed a comprehensive evaluation. First, we retrieved human TF-target interactions for 1,541 TFs using the most established strategies: 1) literature-curated resources; 2) ChIP-seq binding data; 3) TFBSs predictions on human promoters; and 4) reverse-engineered regulons inferred from normal tissue gene expression profiles from the Genotype-Tissue Expression (GTEx)^37^ project (hereafter inferred regulons). We then evaluated to what extent the evidence supporting the TF-target interactions affects TF activity estimations in three different benchmark datasets: two involving gene expression measurements after TF perturbations and one derived from cell-specific essentiality profiles in cancer cell lines. We also investigated the limitations and benefits of the different TF-target datasets and how these relate to several TF properties such as the mode of regulation (MoR) or regulatory effect on their targets, the mode of interaction with the chromatin, the DNA-binding domains and specificity, dimerization or tissue of expression. Finally, we provide general guidelines for the quantification of TF activities across heterogeneous populations of samples together with the retrieved TF regulons as a resource for the community.

## MATERIALS AND METHODS

### TF census and classification

Here we consider a TF as a protein that binds DNA in a sequence-specific manner and regulates the expression level of the target gene^38^. We used the census of human Transcription Factors from TFClass database (v2014) involving 1,541 human TFs^39^, classified according their DNA-binding domain. Moreover, we annotated each TF according to: 1) the mode of interaction with the chromatin (Pioneers, Settlers and Migrants) using the results from *Ehsani et al^40^;* 2) the number of GTEx tissues^37^ where the gene is expressed (i.e. average expression > 2 fpkm); 3) the DNA-binding mode (monomer, homomer or heteromer) that we manually curated from UniProt^41^ (version November 2017) and complemented with the annotation provided in *Lambert el al*^42^; or 4) their DNA-binding specificity, also from *Lambert el al*^42^. Moreover, we used UniProt “Function CC” field to manually classify TF into activators, repressors, activators and repressors or unknown mode of regulation (MoR). TFs annotation is provided as Supplementary Table S1.

### TF-target data sources

#### Literature-curated resources

We downloaded manually curated TF-target relationships from 12 sources (Fantom4^14^, HTRIdb^15^, IntAct^18^, KEGG^17^, ORegAnno^19^, NFIRegulomeDB^13^, PAZAR^12^, TFactS^8^, TFe^16^, TRRD^10^, TRED^11^, TRRUST^20^) and a manual curation of targets from TF-centric papers^43–47^. From Fantom4, we downloaded the “edge.GoldStd_TF.tbl.txt” file. From HTRIdb (v052016), we excluded interactions derived from large-scale experiments such as Chip-Chip and ChIP-seq. From IntAct, we queried all human protein-DNA interactions involving a TF protein and a gene. From KEGG, we used KEGGREST R library to download all *homo sapiens* pathways and retrieve regulatory interactions classified as “GErel”. From ORegAnno, we separated the relationships from PAZAR and NFIRegulomeDB. The remaining relationships were classified as ORegAnno, keeping the interaction sign (i.e. MoR). From TFactS catalogue, we extracted human interactions and separated TFactS-specific interactions from TRRD interactions using the *REF* field. TRED interactions were retrieved via RegNetwork database. From TFe, we downloaded the manually curated targets and the interaction sign. From TRRUST, we extracted all human interactions. When the same TF-target interaction was assigned with more than one sign (*Activation* or *Repression*), we kept the interaction sign with more *PubMed* references. If the same number of references was observed, we prioritized the sign as follows: *Activation* > *Repression* > *Unknown.* Finally, we manually extracted the canonical targets listed in several TF-specific revision papers^43–47^.

#### ChIP-seq interactions

We downloaded the merged ChIP-seq binding peaks provided by ReMap^48^ from ENCODE and other public resources. For each TF, each binding site is assigned to the closest transcription start site (TSSs) using *bedtools closest*^49^. We obtained human TSSs from Gencode version 26^50^ (GRCh38). For all genes that have multiple transcripts, we chose the closest binding site-transcript pair. Therefore, each binding site is assigned to one gene but each gene may have 0 or more binding sites for a given TF. For each binding site-gene pair, we assigned a binding site-gene score between 0 and 1 based on the distance between the binding site and TSS, similar to Ouyang et al^51^: 
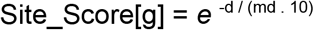
where *md* denotes the median distance between TSS and binding site for each TF and *d* is the distance between each binding site and the TSS of each gene *g*. Therefore, the same score can be assigned a different distance for each TF depending on whether it tends to bind proximally or distally. The score for each TF-target gene assignment is the sum of the scores for all binding sites of that TF assigned to that gene. Finally, for each TF, all TF-target scores are scaled to a value between 1 and 1000. Scores for the top 500 targets of each TF are provided in Supplementary File 1.

#### TF binding sites (TFBSs) predictions in promoters

For each transcript, we scanned human promoter sequences (version GRCh38) for TFBSs. These were defined as the genomic sequence that comprises 1,000 bp upstream (5’ direction) and 200 bp downstream (3’ direction). Mononucleotide PWMs were downloaded from the HOCOMOCO^29^ (v11) and JASPAR^28^ (v2018) repositories. HOCOMOCO-core (excluding low-quality models i.e. “D” category) comprised 402 whereas JASPAR (Vertebrates, non-redundant) comprised 572 PFMs. Prediction of potential TFBSs in the promoter sequences (on both strands) was done using the motif discovery tool FIMO, from the MEME suite (v4.12)^52^ with the default parameters. We selected FIMO predictions with p-value < 0.0001. Duplicated matches (exact binding sites found in different transcripts of the same gene) have been removed.

Next, we annotated the conservation and epigenetic regulatory features of the TFBSs. Base-level PhastCons^53^ and PhyloP^54^ scores were extracted from CellBase database^55^. PhyloP version “hg38.100way.phyloP100way” and PhastCons version “hg38.100way.phastCons” we downloaded in October 2016. PhastCons scores range between 0 and 1 and indicate the posterior probability that the site is in its most-conserved state at that base position. PhyloP score is the −log(p-value) under a null hypothesis of neutral evolution, where positive sign indicates greater conservation. In both cases, binding site-level scores were defined as the 75% percentile of the single base-level scores of the binding region. To call a region conserved, PhastCons and PhyloP scores have to be equal or larger than 0.95 or 3 (corresponding to a p-value threshold = 0.05), respectively. Finally, we aligned the genomic coordinates of the TFBSs with Ensembl regulatory^56^ features (GRCh38.p10) using CellBase to extract the feature ID and type. We considered the TFBSs aligned to “open_chromatin_region” or “ChIP_seq_region” to be a regulatory site.

#### Regulons inferred by reverse-engineering from gene expression

We used GTEx v6 human gene expression data^37^ from control donors to infer transcriptional regulatory networks for healthy tissues. We downloaded gene-level raw counts for 18,737 samples from the ExpressionAtlas^57^. 144 samples with more than 30% of 0 raw counts were discarded. Also, we removed genes with an average log counts per million (CPM) lower than 0. Next, we normalized the data using the TMM method implemented in *edgeR R* package (v3.14.0). Then, we used the *voom* function in *limma* package (v3.28.21)^58^ to obtain fitted log2 counts per million. To account for potential sample batch effects, we downloaded the annotation file GTEx_Data_V6_Annotations_SampleAttributesDS.txt from the gtexportal.org and extracted the isolation batch field “SMNABTCH” and corrected it using *ComBat* function from *sva R* package (v3.20.0)^59^. Samples from cell lines were discarded. Finally, replicates (i.e. samples from the same tissue and donor) were averaged. This data covers 9,407 samples from 30 tissues (histological types).

Next, we used the ARACNe software^32^ (version v1.4) to reverse engineer tissue-specific networks. For each tissue with at least 15 donors, we first pre-calculated ARACNe threshold with a fixed seed with --*calculateThreshold* parameter. Secondly, we ran 100 reproducible bootstraps with a controlled seed and, with the --*consolidate* parameter, derived the tissue-specific network. Finally, we used the *aracne2regulon* function in *viper* R package^2^ (version 1.12.0) to infer the sign of each TF-target interaction (i.e. MoR; activation or inhibition).

We also aggregated the data from all the tissue-specific regulons to infer four consensus regulons, by selecting TF-targets signed interactions appearing in at least 2, 3, 5 and 10 tissues, respectively.

### Benchmark data

Three different benchmark datasets (B1, B2 and B3) were used to evaluate the TF-target resources: B1 and B2 based on gene expression data upon TF perturbation and B3 based on combining basal gene expression with essentiality profiles in cancer cell lines.

#### TF perturbation experiments

##### Benchmark B1

We downloaded microarray expression data corresponding to 189 manually curated TF perturbation experiments in human cell lines from 130 GEO studies. We considered only experiments that fulfil two requirements: 1) provide comparable control and perturbed samples; 2) and have at least two replicates per condition. In each experiment, controls and perturbed samples were manually classified into *positive* (with higher expected activity for the perturbed TF) or *negative samples* (with lower expected activity). Overexpression and knockout-based experiments where the perturbed TF was not differentially expressed were excluded. A total of 94 unique TFs were covered across the collected experiments. When CEL files were available for experiments carried with Affymetrix platforms, we used the functions *ReadAffy* and *rma* from *affy* R package (version 1.50.0) to load and normalize the raw data. Otherwise, we used the expression matrix provided by the authors and applied a quantile normalization for non-normalized data. Finally, for each experiment, we performed a differential expression analysis between the *positive* and the *negative* samples using *limma* R package^58^ (version 3.28.21). Each perturbation experiment was analyzed independently.

##### Benchmark B2

Also, we considered two additional high-throughput shRNA perturbation experiments knocking-down several TFs with no replicates. One in A375 cell line (GSE31534) and another in MCF7 breast cancer cell line (GSE31912)^60^. The corresponding microarray expression datasets were downloaded and normalized as described for the benchmark dataset B1. Normalized expression values were z-transformed to bring the expression from different genes to a common scale. To remove unsuccessful shRNA cases, knocked-down TFs whose expression was in the top 20th percentile (TF-wise) were excluded. 13 and 28 TFs from GSE31534 and GSE31912, respectively, were finally used.

#### Basal cancer cell lines data

##### Benchmark B3

Additionally, we retrieved basal gene expression data from a panel of cancer cell lines that we combined with phenotypic data from two gene shRNA essentiality screens and Copy Number Alterations (CNA) to define likely active and inactive TFs. Specifically, we used gene expression data from our previous publication^4^, that included basal RNA-Seq measurements from three cancer cell line panels: Genomics of Drug Sensitivity in Cancer (GDSC)^61^, Cancer Cell Lines Encyclopedia (CCLE)^62^ and Genentech^63^. Data is available to download from the Expression Atlas^57^ under the accessions E-MTAB-3983, E-MTAB-2770 and E-MTAB-2706, respectively. Regarding the gene essentiality screens, we downloaded the DEMETER scores from Achilles dataset (v2.20.2)^64^ and ATARiS scores from the project DRIVE^65^. A TF was considered to be essential in a cell line (i.e. positive control) if the DEMETER or ATARiS z-scores were < -4 and non-essential (i.e. negative control) if the z-scores were > 4. TFs carrying homozygous deletions were also used as negative controls. CNA for the cell lines were downloaded from the GDSC portal^61^.

### TF-activity scoring methods

We estimated TF activities as a proxy of the expression levels of the targeted genes using the analytic Rank-based Enrichment Analysis (aREA) method from the *viper* R package^2^, a statistical test based on the average ranks of the targets. For perturbation experiments in B1, changes in TF activities for each perturbation were derived from the differential expression signatures via *aREA-msviper* function. For GSE31534 and GSE31912 experiments in B2 and the cancer cell lines in B3, TF activities were derived from the z-transformed expression values via *aREA-viper* function. In both cases we used the default parameters, with the exception of *ges.filter/eset.filter* that was set to FALSE to avoid limiting the expression signatures to the genes represented in the regulons. The Normalized Enrichment Score (NES) was used as a measure of relative TF activity. NES were estimated for each TF in each individual regulon dataset independently (ex. *TP53 regulon from IntAct, TP53 regulon from ReMap*, etc). To avoid confounding effects, self-regulatory interactions were removed. Also, overrepresented targets (regulated by more than 10% of the TFs in the regulon dataset) were discarded. Only TF regulons with at least five targets were tested.

The aREA method takes into account the sign of each TF-target interaction. Here, when the MoR of the TF-target interaction was not defined by the original dataset (i.e. those derived from TFBS predictions, ChIP-seq data and most of the curated databases), we assumed a positive regulatory effect of the TF on the target. However, if the TF is known to be a global repressor (data extracted from Uniprot; Supplementary Table S1), the interactions are assumed to have a negative regulatory effect.

### Comparing regulons performance

To evaluate the performance of each regulon resource, we compared the estimated activities of the TFs in each benchmark dataset. In the context of the perturbation-based benchmark datasets B1 and B2, our assumption is that if the TF regulon defined by a given source (e.g. TP53-IntAct) is accurate, then the experiment perturbing such TF (positive controls) should display lower activity than the rest of perturbations (negative controls) and rank at the top. In the same way, in our benchmark dataset B3 we expect that the sample where a TF is essential (and therefore active; positive controls) should display the highest activity scores while the inactive TFs (negative controls) should take the lowest values. Thus, for each TF regulon under study, we rank the samples according to the TF’s NES. Next, to evaluate the ranking values of the positive and negative controls we performed precision-recall (PR) and receiver operating characteristics (ROC) analyses by the PRROC R package^66^ (version 1.3) and we used the areas under the curves (AUPRC and AUROC) as performance metrics. Since the number of positive and negative controls is unbalanced for the benchmark datasets B1 and B2 (in favour of the negatives), we down-sampled the negatives 100 times to equal the number of positives and took the average metric values.

## RESULTS

### TF-target datasets description and overview

First, we retrieved putative direct transcriptional targets for 1,541 human TFs, as defined by TFClass^39^ (Supplementary Table S1), using different strategies that we grouped according to the evidence type: 1) literature-curated collections from publicly available databases^8,10–20^; 2) ChIP-seq interactions from ReMap^48^; 3) TFBS predictions on gene promoters using TF binding motifs from HOCOMOCO^29^ and JASPAR^28^; and 4) transcriptional regulatory interactions across human tissues inferred from GTEx expression data^37^ using ARACNe^32^ (Figure 1A). From the literature-curated resources, we directly retrieved all the TF-target interactions as indicated in the corresponding databases. For ChIP-seq, we downloaded the binding peaks from ReMap and scored the interaction between each TF and gene according to the distance between the TF binding sites and the genes’ transcription start sites. We evaluated different filtering strategies that consisted of selecting only the top scoring 100, 200, 500 and 1,000 target genes for each TF. For TF binding predictions on promoters, we used FIMO scanning tool with TF motifs from HOCOMOCO and JASPAR, identifying 16 and 13 million binding events (p-value < 0.0001), respectively. Again, we studied different filtering strategies to select the top 100, 200, 500 or 1,000 unique hits and filtering these according to the conservation of the promoter binding sequence or chromatin accessibility annotations from Ensembl. Finally, for the prediction of transcriptional interactions, we ran ARACNe and VIPER on each GTEx tissue independently to build tissue-specific regulatory networks. We also selected interactions identified in at least 2, 3, 5 or 10 tissues to derive consensus inferred TF regulons. See the Methods section for more details. Collectively, our data set contains 1,5 million interactions between 1,399 TFs and 27,976 target genes (Supplementary Table S2).

We then compared the TFs covered per evidence type (Figure 1B). For 101 TFs (6.7%) we retrieved no targets using the mentioned strategies; 638 TFs (42.4%) were covered by a single strategy, being most of these reported by transcriptionally inferred regulons alone (578 TFs; 38.4%). On the opposite side, for 462 TFs (30.7%) we were able to identify targets via three or more lines of evidence. Enrichment analysis revealed that the TFs covered by at least three strategies were expressed across the majority of human tissues (>90% GTEx tissues; Fisher Test, FDR=1.77×10^−6^) and enriched in Cancer Pathways (KEGG; Fisher Test, FDR=2.41×10^−18^ Figure S1). Moreover, these were enriched in Basic Helix-Loop-Helix (bHLH), Basic Leucine Zipper (bZIP), Nuclear Receptors with C4 Zinc Fingers and Tryptophan Cluster factors (Fisher Test; FDR<0.05) while C2H2 Zinc Finger factors are underrepresented (Figure 1C). This last TF class is only covered by transcriptional predictions (Fisher Test; FDR=1.91×10^−12^).

**Figure 1.**
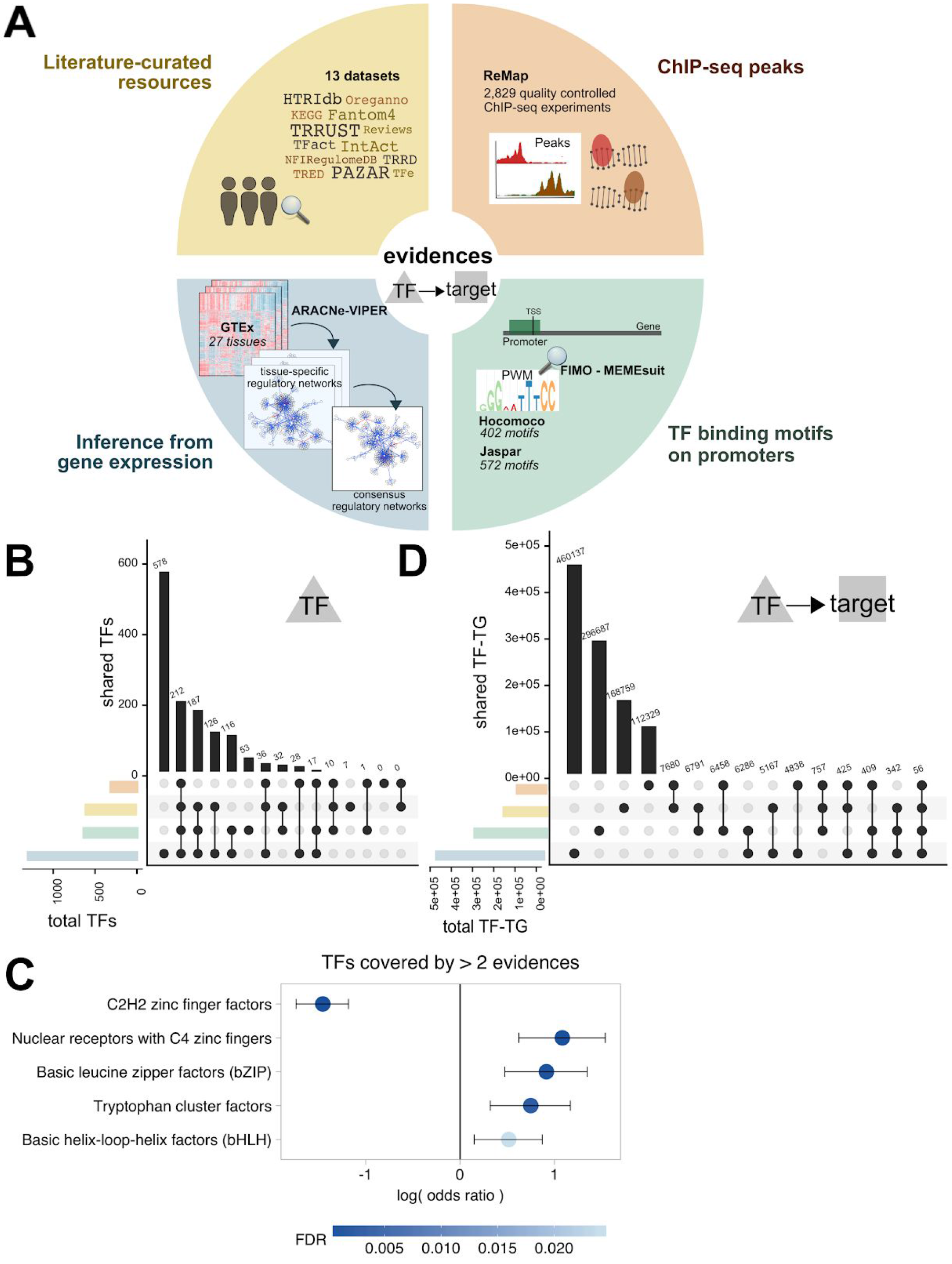
*A) Summary of the resources and strategies used to derive human TF-target interactions, classified according to the evidence level: manually curated resources (yellow), ChIP-seq binding experimental data (orange), prediction of TF binding motifs on gene promoter sequences (green) or inferred from GTEx data (blue). B) TF coverage from each evidence class. C) TF classes (from TFClass) enriched in the TFs covered by more than two lines of evidence. Dots indicate the log odds ratio while error bars indicate the confidence interval. Colors indicate the FDR. D) TF-targets coverage from each evidence class. Note that for regulons inferred from GTEx, only TF-targets > 2 tissues are shown. For TFBSs and ChIP-seq, only top 500 unique hits are shown, p<0.0001.*

We subsequently compared the TF-targets covered across the different strategies (Figure 1D). Notably, the majority of TF-target interactions (96.3%) were supported by a single line of evidence (51.7% where the evidence was a computational prediction). 37,220 (3.4%) are supported by 2, 1,933 (0.2%) by three and only 56 by all four lines of evidence.

### TF-target datasets’ benchmark

Next, we attempted to evaluate the strategies to define TF-target regulons according to their ability to predict changes in TF activities. We reasoned that if the set of targets retrieved for a TF is reliable (i.e. their expression is directly regulated by the TF), the collective expression level of the regulon should be informative of the transcriptional activity of the TF. To determine whether a TF-target dataset provides accurate TF regulons for activity inference, we evaluated the changes in TF activities in three benchmark datasets. First, we manually curated gene expression experiments from GEO including TF knockouts, TF overexpression or TF modulation using a targeted compound. We argued that these experiments are expected to lead to the perturbation of TF activities and, therefore, could be used as benchmark datasets. In total, we collected 189 low-throughput (benchmark dataset B1) and two high-throughput (benchmark dataset B2) perturbation experiments (Figure 2A). Since perturbation experiments are likely to be biased toward well-studied TFs, which may have been thoroughly evaluated in the curated resources, we decided to include a third benchmark dataset (that we called B3) where the positive controls are defined in a data-driven way from a genome-wide analysis. Specifically, for B3 we used two recently published high-throughput gene essentiality screen in cancer cell lines^64,65^ (for which basal gene expression data is available) to identify putatively active TFs. Benchmarks B1, B2 and B3 covered 94, 33 and 135 unique TFs, respectively.

**Figure 2.**
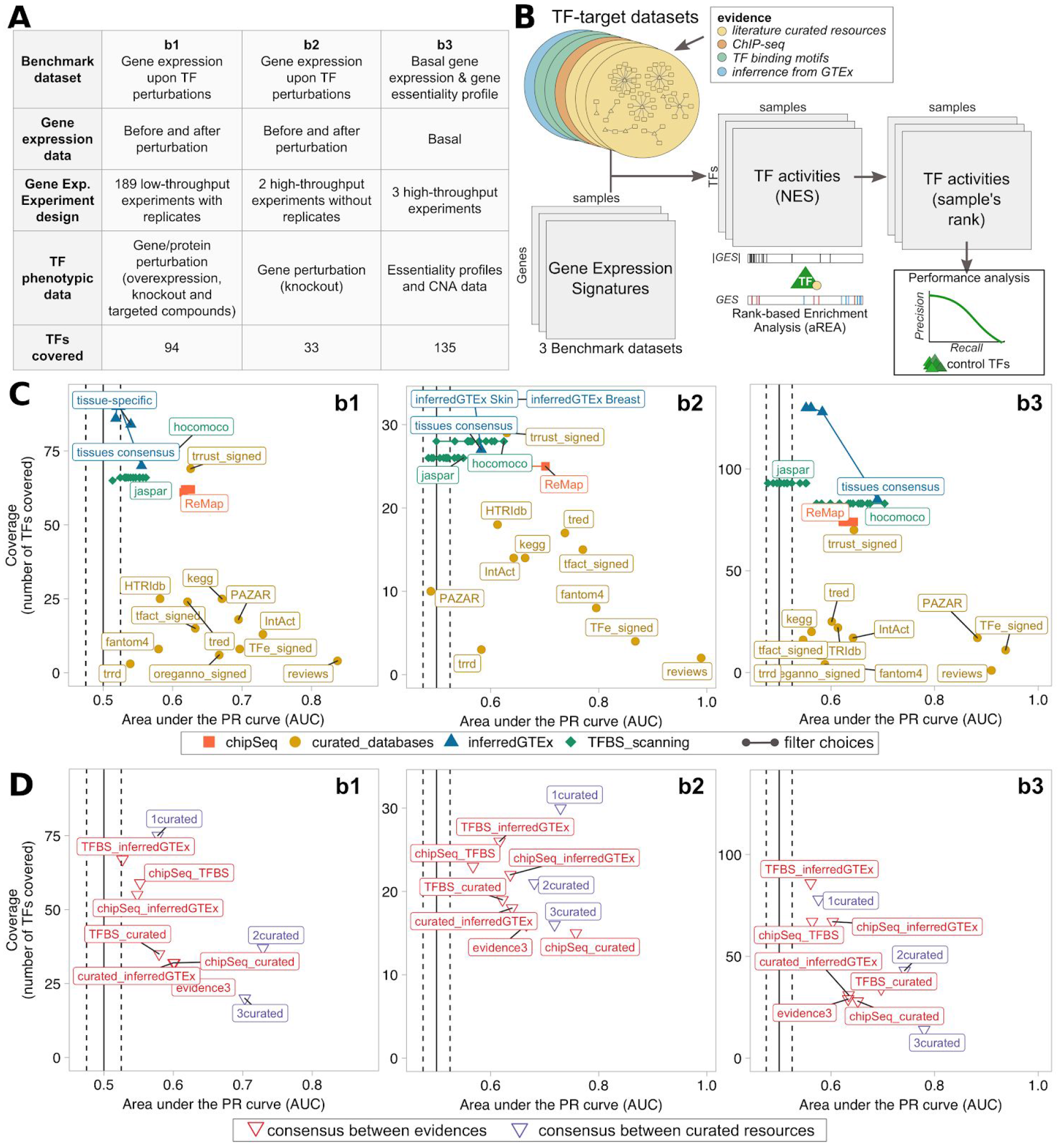
*A) Description of the three benchmark datasets. B) Benchmark analysis scheme. C-D) Performance comparison of the regulons datasets (isolated or in combination, respectively), in terms of TF activity prediction, against the three benchmark datasets. Confidence* vs *coverage plots where X-axis represents the AUPRC from the activity rank’s position of the perturbed/essential TF with respect to the negative controls; and Y-axis represents the number of TF (with ≥ 5 targets) in the benchmark covered by each regulon dataset. Dot colors indicate the evidence type (C, single datasets/evidence) and the nature of the combination (D, combined evidence). For the combined evidence, we considered only TF-target supported by an agreement of two (or three) of any of the four mentioned strategies.*

With the data in hand, the first step in our benchmark strategy (Figure 2B) consisted of estimating TF regulon activities from the gene expression signatures (i.e. expression-level statistic). These gene expression signatures are derived differently across the benchmark datasets due to their differences in the experimental design. For the B1 dataset, since it contains replicates for control and perturbed samples (knockout, overexpression, etc), we defined the gene expression signature of the perturbation as the differential expression between the positive (with expected higher TF activity) and the negative samples (with expected lower TF activity). In contrast, B2 dataset does not contain perturbation replicates. Here, we normalized the gene expression profiles using z-score transformation to derive the expression-level statistic, which quantifies whether a gene in a sample is more or less expressed in the context of the B2 population distribution. B3 does not contain perturbation data, only basal gene expression from unperturbed samples. Here, the gene expression signatures are derived as for B2. With the gene expression signatures in hand, our second step applied the analytic Rank-based Enrichment Analysis (aREA) method to infer the TF regulon normalized enrichment scores (NES) that we used as estimates of TF activities. Finally, NES values were used to rank, for each TF regulon, the experiments or samples within each benchmark dataset. The estimated TF activities were then evaluated against our benchmarks using a Precision-Recall analysis (see Methods), we compared the ranked TF activities estimated for our positive samples (i.e. the perturbed TF-sample pairs in B1 and B2 and the essential TF-sample pairs in B3) against the negative samples (i.e. samples with a different perturbed TF in B1 and B2 and the inactive TF-sample pairs in B3) and quantified the AUPRC and AUROC for each TF-target dataset.

The predicted TF activities derived from most of the resources performed better than random (AUPRC > 0.5, Figure 2C, S2 and AUROC > 0.5 Figure S3). TF regulons manually reviewed by experts, such as those listed in TF-centric review papers or in the TFe resource, reached the highest accuracy levels. In contrast, TF regulons determined *in silico* from TF binding motifs or inferred from data reached higher TFs coverage but lower accuracy. ChIP-seq derived regulons display intermediate performances and coverage. Overall, there is an inverse relationship between the coverage and the accuracy across regulon datasets.

We also explored how the performance of the ChIP-seq and *in silico* methods depend on different filters or parameters which alter the resulted regulons (Figure S2 and S3 for AUPRC and AUROC, respectively), observing differences across the three different benchmark datasets. We reasoned that these divergences may likely be due to, on the one hand, the different conditions and cell models used and, on the other hand, the differences in number and type of TFs covered in each experiment. Still, we observed a global tendency when comparing some filtering features. For example, we evaluated different regulon size cutoffs (top 100, 200, 500 and 1,000 targets) for the ChIP-seq and TF binding predictions on promoters (Figure S2A-B and S3A-B) observing that intermediate cutoffs reach higher AUC values overall. The use of chromatin accessibility information to further filter TF binding hits on promoters decreased their performance while the inclusion of sequence-based conservation reached similar or better performances (Figure S2C and S3C). TF binding motifs from HOCOMOCO performed generally better than the ones from JASPAR, considering only TFs covered by both resources (Figure S2B-C and and S3B-C), likely because the former is based on experiments only from human while the latter also from other Vertebrates. Lastly, for the overlapping TF regulons inferred from GTEx, we observed that tissue-specific interactions (i.e. interactions detected in the GTEx tissue matching the tissue lineage of the samples in the perturbations) performed similar to consensus regulons (i.e. interactions detected in more than one GTEx tissue), where consensus TF-target interactions observed in more tissues performing better (Figure S2D and S3D).

We also asked whether the inclusion of the mode of regulation (MoR) provided for some TF-target interactions in some literature-curated databases had an impact on the prediction of the TF activities. Comparison of signed and unsigned version of the same regulons revealed similar performances, with the exception of TRRUST database (Figure S2E and S3E). The signed version of TRRUST regulons improved significantly the TF activity predictions (Wilcoxon-test; p<0.05) for all three benchmark datasets.

Finally, we selected the TF-target interactions supported by more than one line of evidence and recomputed the activities for the resulting TF regulons across the benchmark datasets (Figure 2D). Globally, regulons containing TF-targets supported by at least two literature-curated resources displayed the best performances across the three benchmark datasets. In contrast, regulons built by intersecting *in silico* predictions perform poorly, with no improvement compared to the use of regulons uniquely derived from each single strategy alone. Interactions supported by at least one literature-curated resource and ChIP-seq, or supported by any three lines of evidence showed an intermediate performance.

### TF properties affecting inference of regulon activities

Not all the TFs function in the same way or are regulated by the same mechanisms. TFs may differ in their MoR, the way in which they interact with the chromatin, the conditions upon which they are expressed or the regulation by post-translational modification or the interaction with cofactors. In order to characterize if any of these properties could affect the power to infer accurate TF activities, we annotated, when possible, our list of human TFs according to their global regulatory effect on the targets (i.e. activators, repressors or dual) in Uniprot^41^, if these operate as complexes (i.e. heteromers or homomers) according to Uniprot^41^ or *Lamber et al*^42^, the DNA-binding specificity^42^, the mode of interaction with the chromatin (Pioneers, Settlers and Migrants)^40^ and their classification based on their DNA-binding domains from TFClass (Figure 3A). Additionally, we classified the TFs as tissue-specific or widely expressed if their transcripts were detected in less than 10% or more than 90% of the tissues in GTEx (Supplementary Table S1; see methods section for details).

**Figure 3.**
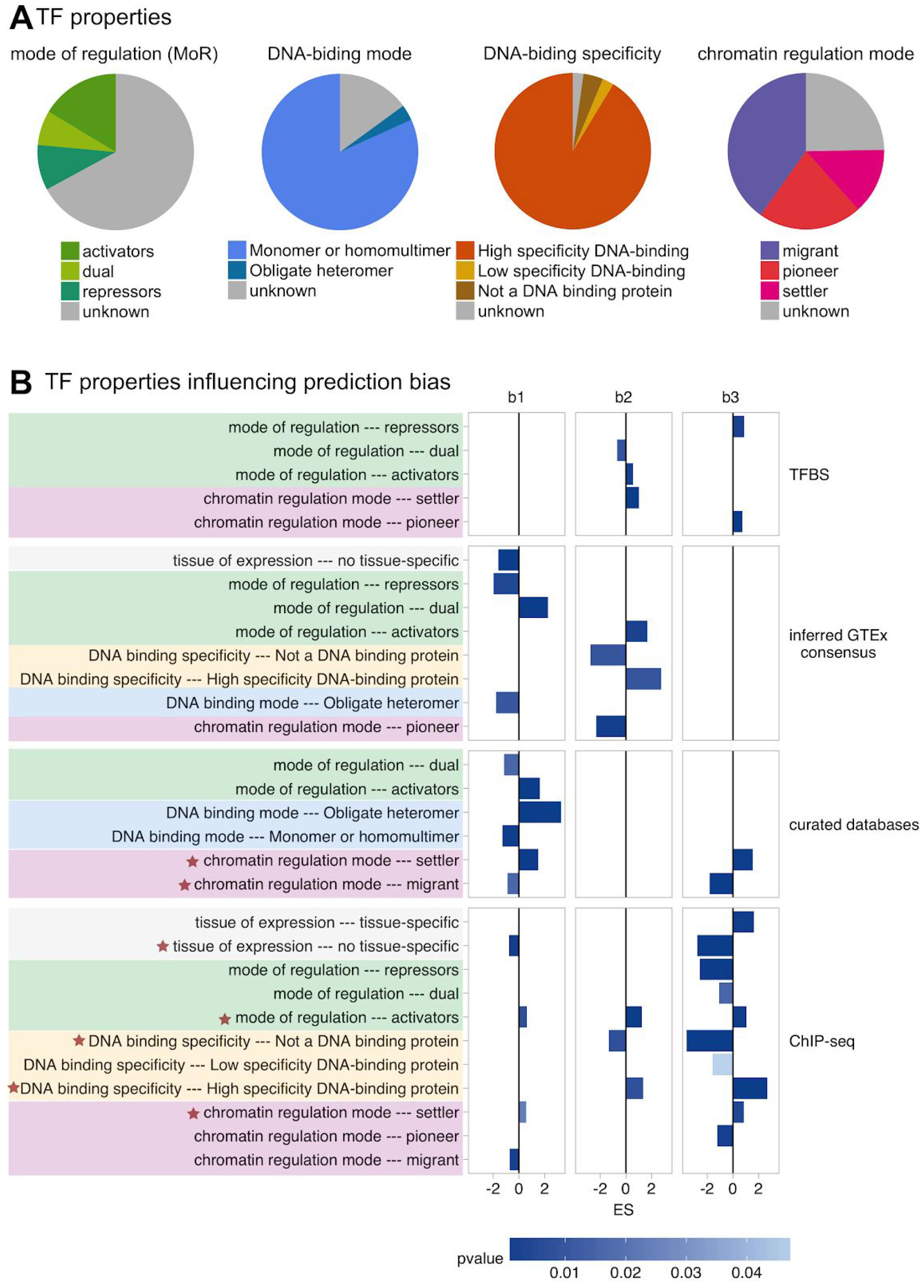
*TF properties biasing the inference of TF activities across the TF regulon datasets. A) Overview of the TF properties annotated for the 1,541 human TFs under study. B) TF properties enriched (FDR<0.01) in the benchmark results B1, B2 and B3. Bar length in proportional to the Enrichment Score (ES) while color represents the significance strength (p-value). Properties enriched in more than one dataset are labeled with an asterisk.*

GSEA analysis^67^ of the benchmark results revealed that indeed certain TF properties associate with better and worse TF activity predictions in the context of certain benchmarks and TF regulon datasets (Figure 3B). For example, when evaluating the DNA-binding specificity of the TFs, those classified as low specificity are worse predicted than those with higher DNA-binding specificity, using ChIP-seq and expression-inferred regulons. Focusing on the MoR of their targets, TFs that are expected to act only as activators or repressors are better predicted than TFs with dual MoR, using regulons from ChIP-seq, TFBS and literature-curated sources. This is probably caused by the incomplete knowledge on the MoR of TF-target interactions in these datasets. In contrast, TFs with a dual MoR are better predicted using consensus regulons inferred from GTEx, likely due to the inference of the TF-target interaction sign from the gene expression patterns. Also, we observed that the properties related to the regulation of the TF function strongly impact the performance of the predictions. For example, TFs binding the DNA as heteromers were poorly predicted using regulons inferred from expression data, likely due to the fact that the inference approach does not consider interactions between TFs in their model. Similarly, when studying the impact of DNA accessibility, our results showed that ‘settler TFs’, which bind to all the accessible DNA sites matching their DNA-binding motifs, display better predictions across all regulon types except for those inferred from gene expression. In contrast, TFs with more complex interactions with the chromatin and cofactors, called ‘migrant TFs’ (binding only to part of the accessible DNA sites matching their DNA-binding motifs), are worse predicted using ChIP-seq and literature-curated regulons. Finally, we also observed that TFs ubiquitously expressed across GTEx tissues, likely involved in a wider range of processes and intricate regulatory mechanisms^38^, are worse predicted than TFs with tissue-specific expression (1 or two GTEx tissues). Taking together, these results indicate that activity predictions are less accurate for TFs under complex molecular control.

### An integrated resource of scored TF-target interactions

With the aim of providing a comprehensive resource of regulatory TF-target interactions, we have integrated all the collected interactions from the four lines of evidence and assigned a confidence score to each one based on the benchmark results. This resource incorporates: 1) all the interactions derived from the literature-curated resources; 2) the top 500 scoring interactions per TF identified from the collection of ChIP-seq experiments from ReMap; 3) the top 500 interactions per TF identified by scanning the human gene promoters with JASPAR and HOCOMOCO motifs (p < 0.001); and 4) the regulons inferred by ARACNe in at least three GTEx tissues. This resource, available in *OmniPath* (www.omnipathdb.org)^68^, and at https://saezlab.github.io/DoRothEA/, comprises 1,077,121 TF-target candidate regulatory interactions between 1,403 TFs and 26,984 targets.

Here, for each TF-target interaction we assigned a confidence score based on the observations from our benchmark. The score comprises five categories, ranging from A (highest confidence) to E (lowest confidence). The scoring criteria is described in Figure 4A. Briefly, interactions that are supported by all the four lines of evidence, manually curated by experts in specific reviews or supported both in at least two curated resource are considered to be highly reliable and were assigned an A score. Scores B-D are reserved for curated and/or ChIP-seq interactions with different levels of additional evidence. Finally, E score is used for interactions that are uniquely supported by computational predictions. Figure 4B describes the TFs and interactions coverage per scoring category.

Finally, we used the scored regulons to characterize the TF activities our benchmark datasets. As expected, TF activities derived from A-B scored target regulons perform notably better than activities estimated using E-scored targets in the three benchmark datasets.

**Figure 4.**
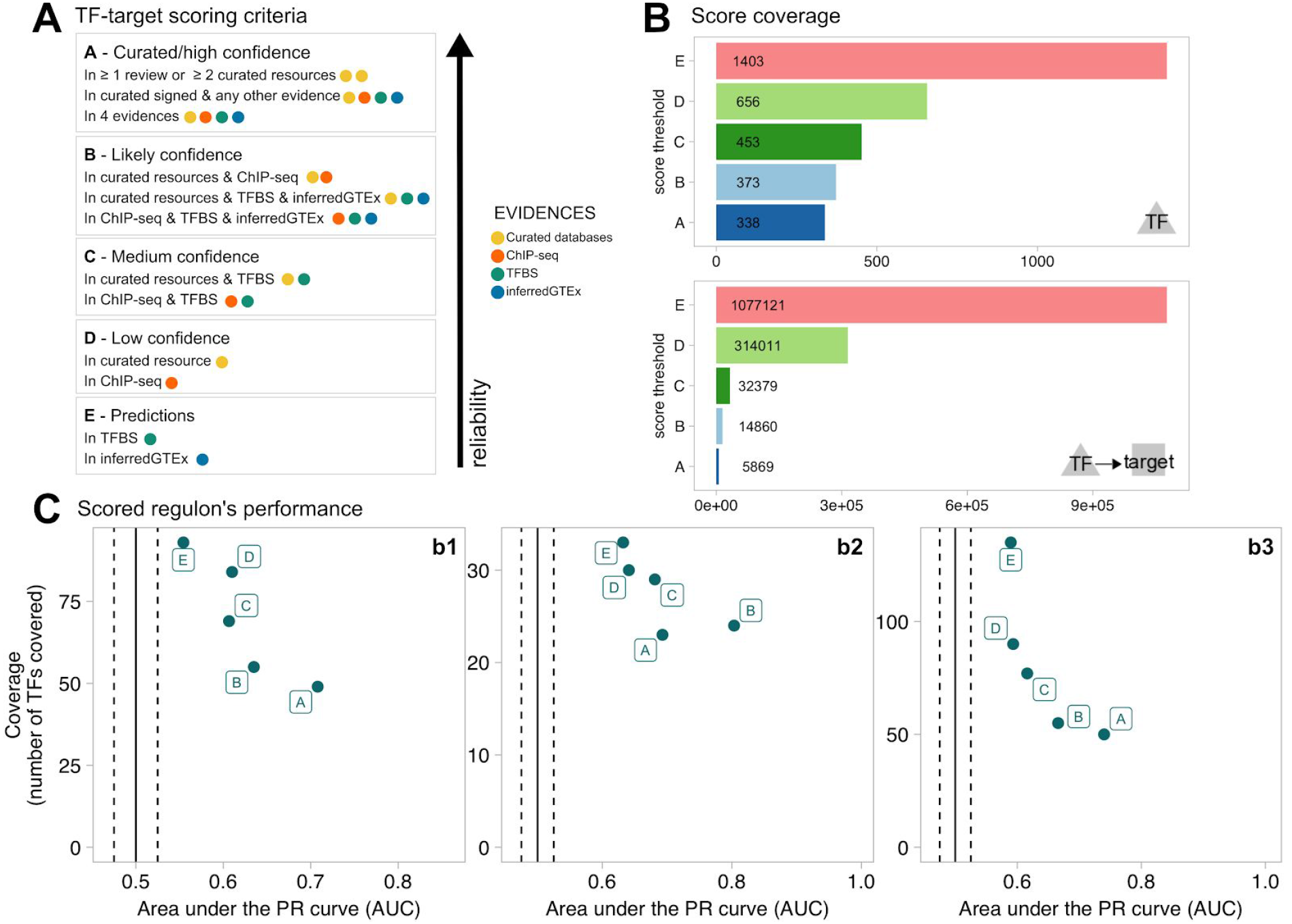
*Scoring TF-target interactions from different evidence. A) Scoring scheme. B) TF and TF-target interaction coverage per score cutoff. C) Performance of scored regulons in B1, B2 and B3 benchmark datasets.*

## DISCUSSION

Inference of TF activities from the expression levels of their putative targets is becoming a widespread tool to extract functional insight from transcriptomic data^2–7,69^. Although several strategies exist to define the TF’ targets (i.e. TF regulon), their potential to predict changes in TF activities has not yet been systematically compared. To employ these data conveniently, a critical evaluation of their reliability (i.e. quality), coverage (i.e. quantity) and complementarities is needed. Here we evaluated the impact of the four major types of evidence supporting the human TF-target interactions in the estimation of TF activities: literature-curated repositories, ChIP-seq binding data, *in silico* predictions of TFBSs on gene promoters and inference by reverse-engineering from large gene expression datasets.

Overall, we observed that for almost half of the TFs, only one of the four strategies report targets. More remarkably, there is low overlap at the TF-target interactions-level, were the great majority are supported by a single line of evidence. Therefore, it seems that to have a complete picture of the human TF-target regulatory map, integration of different strategies is essential.

Comparing the quality of TF-target interaction datasets is complex due to their relatively low coverage and overlap. More importantly, these need to be benchmarked against a reliable and comprehensive reference set. Up to our knowledge, there is no systematic experiment measuring directly the protein-level activities of hundreds of human TFs in several conditions. Under these limitations, in order to compare the different TF-target resources and detection strategies, we collected three alternative benchmark datasets where changes in TF activities are assumed indirectly. In general, although there is a tendency (curated regulons are better than ChIP-seq, which in turn are better than predicted ones), we noticed differences in the performance of each TF-target strategy across the benchmark datasets. This can be due to various technical and biological factors. Among the technical factors, we note the low coverage and overlap of the TFs included in each benchmark experiment, as well as the different experimental conditions and assumptions used to derive these control datasets. In fact, in the perturbation-based benchmark datasets B1 and B2, some TFs in the negative controls (i.e. not directly perturbed TFs) can be indirectly affected and, therefore, not represent true negatives. These could involve, for instance, TFs co-regulated by the perturbed TFs. Moreover, some of the TFs in the positive set (i.e. perturbed TFs) may not be effectively modulated by the perturbation strategy used (overexpression or knock-out). For example, the experiments overexpressing a TF gene may not translate into an efficient activation of the coded protein if the regulatory elements (e.g. post-translational modifications) or external stimulus (e.g. viral infection) needed for such activation are not present. In fact, when comparing TFs with different regulatory properties, performances are generally challenged when predicting activities for TFs with dual MoR (that can act as activators or repressors of their targets) or under complex molecular control, such as those working as heterodimers, interacting with cofactors or other chromatin regulators. In these cases, the combination of complementary strategies to define TF-targets can enhance the TF activity predictions. Additionally, other biological factors influence the performance of the benchmark such as the modulation of TFs other than the perturbed ones due to cooperativity, feedback regulation or redundancy mechanisms in the regulation of transcriptional programs, which used to buffer loss-of-function perturbations or result inappropriate activation of specific TFs. Taking all together, we hypothesize that the limitations of the benchmark datasets are masking and underestimating the potential of all these TF-target resources.

Still, our results highlight the importance of the literature-curated regulons to infer accurate TF activities, with the highest precision achieved for interactions supported by more than one resource or expert’s review. However, their systematic use is limited by the low coverage, what requires the integration of multiple resources. Among literature-curated resources, noteworthy is to mention the TRRUST database, which displays the best balance between regulon performance and coverage, the latter likely boosted by the systematic sentence-based text mining search of regulatory interactions prior to manual curation and the incorporation of the MoR. Focusing on high-throughput strategies, ChIP-seq binding regulons displayed the highest performance with a coverage comparable to that of TF binding motifs on gene promoters. In general, *in silico* predictions based on TFBSs on gene promoters or inferred from GTEx’ displayed the lowest performance among all strategies. This is likely due to the higher ratio of false positives intrinsic to the technical biases of each strategy^70,71^. Interestingly, tissue-specific regulons inferred from GTEx data perform worse than inferred consensus regulons (i.e. selecting TF-target interactions detected in several tissues). A reason may be the inherent differences between the gene expression regulatory programs in the samples used to infer the regulatory networks (normal tissues from GTEx) and in the samples in the benchmark (cell lines, mostly cancer-derived).

When combining TF-target interactions supported by both curated databases and ChIP-seq or any three lines of evidence, the performance increases with respect to the use of interactions only reported by literature-curated resources, suggesting that these can be further refined integrating other lines of evidence. In contrast, regulons detected by the two *in silico* prediction methods did not improve the performance with respect to the use of these alone.

With these observations in mind, we have integrated 1,077,121 TF-target interactions derived from the mentioned strategies that we have accompanied with a confidence score. Up to our knowledge, this is the largest collection of human TF-targets interactions from heterogeneous sources and strategies. This integrated resource can be of broad applicability for approaches requiring the inference of the regulatory activity of TFs, where researchers can decide the level of confidence and coverage they want to use in their studies.

The use of regulons to estimate TF activities has many applications, and can be particularly powerful to extract signal robustly from noisy or low coverage expression data such as in the case of single-cell RNA data^4,9^. In addition TF activities can be linked to upstream signaling pathways. Pathway activities are often inferred by the transcription levels of their members, ignoring the hard-to-measure post-transcriptional and post-translational regulation. However, considering gene expression as a downstream effect of pathway activity instead leads to more accurate estimations^72,73^. Further on, TF activities can be used to infer the activity of upstream proteins using knowledge on pathways and how they impinge on TFs^74–76^. The resources and confidence estimates we propose will support the development of such methods. More in general, we expect the presented comparative assessment of the TF regulon resources to contribute to the establishment of guidelines for the quantification of human TF activities.

## ACKNOWLEDGEMENTS

We thank Christian Holland and Anthony Mathelier for useful feedback on the manuscript.

